# A computational method for immune repertoire mining that identifies novel binders from different clonotypes, demonstrated by identifying anti-Pertussis toxoid antibodies

**DOI:** 10.1101/2020.06.02.121129

**Authors:** Eve Richardson, Jacob D. Galson, Paul Kellam, Dominic F. Kelly, Sarah E. Smith, Anne Palser, Simon Watson, Charlotte M. Deane

**Affiliations:** Department of Statistics, University of Oxford, UK; Alchemab Therapeutics Ltd, London, UK; University Children’s Hospital, University of Zurich, Switzerland; Kymab Ltd, Cambridge, UK; Department of Infectious Diseases, Faculty of Medicine, Imperial College London, UK; Department of Paediatrics, University of Oxford, UK; Oxford University Hospitals NHS Foundation Trust, Oxford, UK

## Abstract

Due to their shared genetic history, antibodies from the same clonotype often bind to the same epitope. This knowledge is used in immune repertoire mining, where known binders are used to search bulk sequencing repertoires to identify new binders. However current computational methods cannot identify epitope convergence between antibodies from different clonotypes, limiting the sequence diversity of antigen-specific antibodies which can be identified. We describe how the antibody binding site, the paratope, can be used to cluster antibodies with common antigen reactivity from different clonotypes. Our method, paratyping, uses the predicted paratope to identify these novel cross clonotype matches. We experimentally validated our predictions on a Pertussis toxoid dataset. Our results show that even the simplest abstraction of the antibody binding site, using only the length of the loops involved and predicted binding residues, is sufficient to group antigen-specific antibodies and provide additional information to conventional clonotype analysis.

## 1 Introduction

Next-generation immune repertoire sequencing (Ig-seq or rep-seq [1, 2]) is providing us with comprehensive information about adaptive immune repertoires across individuals [3] and immune states [4]. Progress has been made in the task of interrogating the vast diversity of B-cell receptor (BCR) repertoires, primarily through the analysis of predicted clonal relationships inferred via clonotyping [5]. Ig-seq and associated clonal analysis are finding increasing importance in antibody discovery both as a method of identification of putative antigen-specific antibodies [6–8] and more recently as a method of lead antibody optimization through repertoire mining [9]. The identification of antibodies which are predicted to bind to the same site (epitope) is a key component of BCR repertoire analysis and antibody discovery.

The starting point for most BCR repertoire analysis is the reduction of thousands or millions of BCRs into orders of magnitude fewer clonotypes [5]. A clonotype is defined as a group of antibody sequences which derive from a common progenitor B cell [10]. During B cell development, the variable (V), diversity (D) and joining (J) gene segments encoding the variable domain of the antibody heavy chain undergo recombination [11]. A requirement for two sequences to be predicted to share the same clonotype is therefore common V- and J-germline gene assignment [5]. The D gene is not usually included in standard clonotype definitions due to difficulty in its assignment [12]. The variable domain of the antibody heavy chain consists of the framework regions and hypervariable complementarity-determining regions (CDRs). CDRHs 1 and 2 are encoded by the V gene while the region spanning the recombined V, D and J segments corresponds to the third and most diverse loop on the antibody heavy chain, the CDRH3. The processes of junctional diversification (the insertion of palindromic and random nucleotides at the junction between the V, D and J genes) during recombination as well as somatic hypermutation [11] during affinity maturation further contribute to the diversity of the CDRH3. Sequence identity in the CDRH3 is therefore included as a marker of shared origin in most clonotyping tools [5]. The nucleotide or amino acid sequence identity across the CDRH3 required for two sequences to be considered in the same clonotype varies across studies - in studies performing clonotyping with length-normalised amino acid sequence identity thresholds, sequence identity thresholds vary between 80% and 100% [5]. Published clonotyping methods use only heavy chain information which is sufficient to capture most clonal relationships [13].

A number of publicly available, well-supported pipelines have made clonotype analysis standard practice [5, 10]. This has permitted large advances in the practical utility of Ig-seq data [14, 15]. Clinically, it has found use in tracking minimal residual disease in blood cancers [16], monitoring vaccination responses [17–19] and providing mechanistic insights into immune-mediated diseases [4, 20–22]. Clonotyping has also proven useful in antibody discovery as a means of selecting candidate sequences for expression as monoclonal antibodies [6–8] and recently as a method of lead antibody optimisation via repertoire mining [9].

Antibodies within the same clonotype are likely to target a common epitope [5, 10]. The majority of antibodies binding to the same epitope in antibody-antigen complex structures in the Structural Antibody Database (SAbDab, a database of experimentally-solved antibody and antibody-antigen complex structures) have highly similar CDRH3s [23]. However, it has also been observed that multiple clonotypes may converge on the same epitope. For example, Scheid and colleagues identified clonotypes from distinct IGHV families converging on the CD4 binding site in gp120 [24]. Separate clonotypes have also been observed to bind to overlapping epitopes on the hemagglutinin stem [25] or globular head, and on multiple epitopes on the Ebola virus glycoprotein [26]. Wong and colleagues identified 190 pairs of antibodies with sub-80% CDRH3 amino acid identity binding to the same epitope within SAbDab [23, 27]. This convergence between clonotypes offers the potential to improve our understanding of the functional landscape of BCR repertoires; large-scale functional convergence between lineages could, for example, explain the apparent scarcity of public clonotypes [28]. This hypothesis is supported by evidence that while clonotypes are infrequently shared between individuals, the range of antibody structures that these clonotypes generate is more similar between individuals [29]. In the context of antibody discovery, being able to identify binders to the same epitope from different clonotypes would aid in optimisation of developability or binding affinity, by allowing hopping between germline scaffolds.

In antibody discovery, clonotyping is used to search for clonal relatives of lead antibodies in bulk Ig-seq data sets in order to identify antibodies which target the same epitope but which have either an increased affinity or a superior developability profile. This process is referred to as “immune repertoire mining". Hsiao and colleagues performed clonotyping on a set of bulk repertoires and used the resultant clonotypes for hit expansion against two targets [9]. They achieved greater than an order of magnitude improvement in affinity for both targets and between 48% and 100% of tested heavy chain variants retained target-binding [9]. This suggests that sampling within a clonotype can be highly effective as a means of repertoire mining. However, the method does not allow the identification of binders to the same epitope which derive from different clonotypes, which currently limits the sequence distinctness of novel binders which can be recovered from immune repertoires.

In this paper, we describe a new method to identify functional convergence of antibody sequences that is germline-independent and that considers only the binding site of the antibody sequences, the paratope. We call this approach “paratyping". We show how paratyping allows clustering of antigenspecific sequences from different clonotypes, and is a rapid, structurally intuitive way of clustering functionally-related antibodies. Paratyping simplifies the complex phenomenon of antibody-antigen interaction into sets of shared residues. Learning the complexities of antibody-antigen interactions as part of a predictive model of antigen binding has been achieved in the case of the antigen-interaction of one therapeutic monoclonal antibody [30]. However, such approaches rely on a large (on the order of 10^4^) library of experimentally validated binding and non-binding variants. Paratyping removes the need for large amounts of training data and is generalisable across antigens.

We first show the rationale for our method, paratyping, using the structures of a pair of antibodies from different clonotypes that bind to the same epitope. Paratyping is then applied to a single-cell data set of sequences raised against Pertussis toxoid (PTx) in a transgenic mouse platform where it is as accurate as clonotyping but identifies different binders. We then perform a prospective experimental test of the method by expressing as monoclonal antibodies and experimentally testing predicted PTx-binding and non-binding antibodies mined from a set of non-enriched bulk heavy chain sequencing repertoires. Our experimental test demonstrates that paratyping identifies PTx-binding antibodies from different clonotypes to our original hits. This expands the sequence space available through repertoire mining and permits favourable shifts in in-silico developability metrics. Of particular advantage is paratyping’s ability to prediction common antigen reactivity of antibodies from different V and J gene backgrounds, which has implications for large-scale repertoire analysis.

## 2 Results

### 2.1 Epitope convergence can be identified at the level of paratope residues

Antibodies from different clonotypes have been observed to converge on the same binding site [23–26]. We hypothesise that these functionally convergent antibodies may use the same paratope for interaction, and examined the Structural Antibody Database (SAbDab) [27] for evidence of this. We define the epitope and paratope as those residues with any atom within 4.5 Å of any residue in the cognate antibody or antigen respectively.

Figure 1 shows an example of a pair of antibody-antigen complex structures where antibodies binding to the same epitope derive from different clonotypes but have very similar paratopes. The two monoclonal antibodies 4C2 (PDB ID: 5do2) and D12 (PDB ID: 4zpv) bind to the receptor binding domain of the MERS-CoV spike protein at the same epitope (94.7% of epitope residues are shared across the pair of structures). The antibodies come from different clonotypes with both low CDRH3 identity (66.7%) and different germline origins (V5-6-4/J2 vs V5-9-1/J4). However, 4C2 and D12 use largely the same paratope residues to achieve this epitope convergence, with 81.3% of paratope residues conserved across the structures. Another example of epitope convergence between antibodies with differing J genes and sub-80% CDRH3 identity can be seen in the anti-lysozyme antibodies HyHel-10 and HyHel-63 (see Supplementary Figure 1).

**Figure 1:**
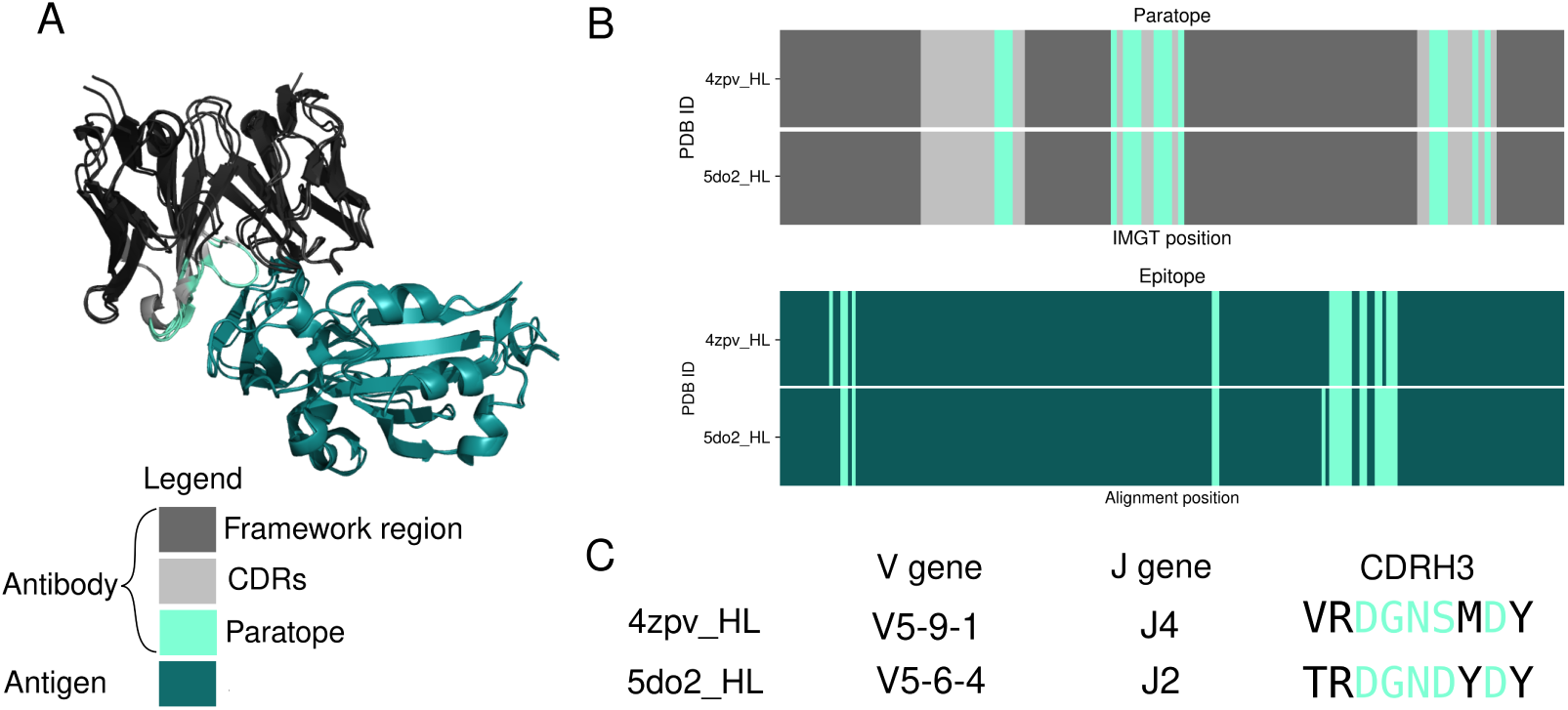
An example of two antibodies that bind to the same epitope but derive from different clono-types. The murine 4C2 (PDB ID: 5do2) and D12 (PDB ID: 4zpv) anti-MERS-CoV antibodies target the same residues on the receptor binding domain of the spike protein, despite being derived from different V and J genes and displaying CDRH3 amino acid identity (66.7%) below the standard clonotyping definition (80% - 100%) [5] (C). The antibodies use over 80% of the same paratope residues (B) to achieve this functional convergence.

### 2.2 Paratyping and clonotyping successfully cluster PTx binders in a single-cell dataset

In a standard implementation, antibodies with the same V and J genes, CDRH3s of the same length, and above a threshold level of amino acid identity across the CDRH3 are considered to be in the same clonotype [5]. In our novel method, paratyping, antibodies with the same length CDRs and above a threshold level of sequence identity across the predicted paratope residues are grouped into the same paratype.

To test the ability of paratyping and clonotyping to group antibodies that target the same epitope we performed a test in a single-cell (paired VH/VL) data set of 1290 antibodies isolated from genetically engineered mice that have a full set of human immunoglobulin variable region genes [31] immunised with Pertussis toxoid (PTx). Although we had pairing information within the single-cell data set, given that the majority of repertoire sequencing data to date is heavy-chain only [32], we demonstrate the method using only the heavy chain information. The sequences were annotated with a PTx-binding (364) or non-binding label (926) (as per the “Single-cell data set” section of Methods), and we used paratyping (our new method) and clonotyping (the conventional approach) to identify PTx-binding sequences.

For each of the 364 PTx-binders in turn, we mimicked a repertoire mining experiment by using paratyping or clonotyping to identify binders amongst the remaining 1289 sequences (one-vs-all cross-validation). Each of the PTx-binders is referred to as a “probe” antibody; sequences that are within the same paratype or clonotype as the probe are predicted to bind PTx. The precision and recall of the two methods (calculated over the aggregate of predictions) for repertoire mining are comparable (Figure 2, Table 1). The precision and recall using clonotyping and paratyping with varying CDRH3 sequence identity or paratope sequence identity thresholds, respectively, are shown in Figure 2A and 2B. The methods require different sequence identity thresholds for optimum performance but have similar precision-recall profiles.

**Table 1:** Precision-recall values for prediction of PTx-binding according to paratyping and clonotyping at the optimal thresholds of 75% and 72% respectively. Sequence identity is calculated across the predicted paratope for paratyping and across the CDRH3 for clonotyping. The methods behave comparably over the full precision-recall curve. These thresholds were then used for the prospective repertoire mining experiment.

| Method | Sequence identity threshold | Precision | Recall |
| --- | --- | --- | --- |
| **Paratyping** | 75% | 84% | 76% |
| **Clonotyping** | 72% | 83% | 79% |

**Figure 2:**
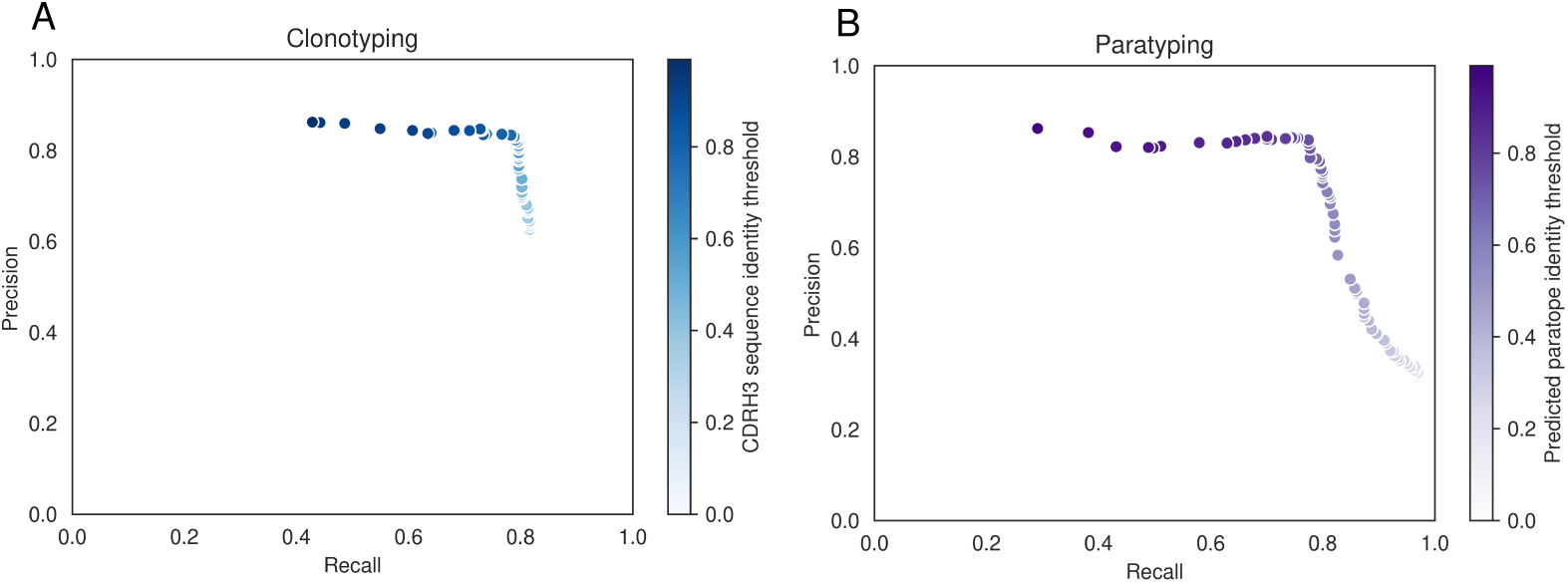
Precision-recall curves for clonotyping (A) and paratyping (B) in the task of predicting PTx binding. Precision and recall values are calculated over a range of CDRH3 sequence identity and predicted paratope identity thresholds respectively between 0.0 and 100.0% identity. The precision-recall profiles are similar.

For clonotyping, the optimal heavy-chain only threshold is 72% CDRH3 amino acid identity. For paratyping, the optimum occurs at 75% paratope identity. At these optimal thresholds, clonotyping recovers binders with 83% precision and 79% recall (meaning that 21% of the binders in this data set are not related by clonotype to any other). Paratyping recovers binders with a precision of 84% and a recall of 76% (meaning that 24% of binders in this data set have distinct paratopes). We expect that this would be an overestimation of performance in the bulk data set due to the enrichment of PTx-binders created through antigen-specific sorting.

For each probe, a prediction can be made by both paratyping and clonotyping (hereafter labelled a “Both” prediction), paratyping alone (labelled “Paratype-only”) or clonotyping alone (labelled “Clonotype-only”). Out of all of these predictions, 76.6% are made by both methods with 89 “paratype-only” predictions (precision: 76%) and 345 “clonotype-only” predictions (precision: 84%). Paratyping and clonotyping make a number of method-exclusive predictions, as shown in Figure 3, where the probe antibodies with the largest number of “paratype-only” and “clonotype-only” predictions are shown in dendrograms with these predictions. Figure 3 also shows the variability of the method’s performance across probes, with the probe antibodies yielding the lowest and highest precision shown. One probe identified 20 other PTx-binding antibodies with 100% precision, while another had just 33% precision. The heterogeneity in predictions across probe emphasises the utility of either method when only a small number of binders are known, despite the similar behaviour of the methods over the aggregate of probe antibodies

**Figure 3:**
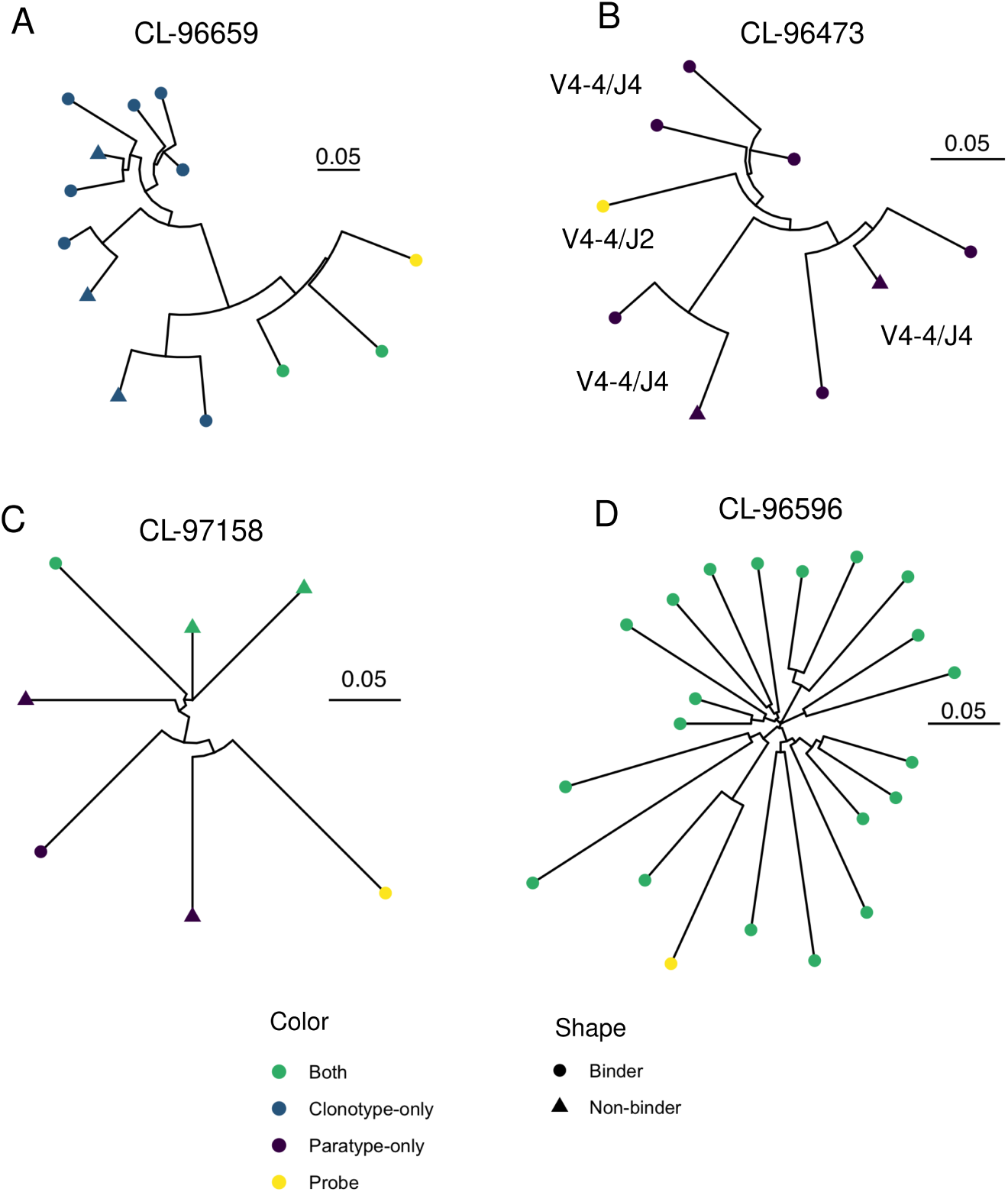
Representative dendrograms from the single-cell repertoire mining experiment. Probe sequences (yellow) are known PTx binding sequences. Other sequences which are in the same paratype or clonotype are predicted to also bind PTx. These predicted PTx binding sequences are coloured according to whether they are identified by both paratyping and clonotyping (“Both”), paratyping but not clonotyping (“Paratype-only”) or clonotyping but not paratyping (“Clonotype-only”). Circular leaves represent true PTx binding antibodies (i.e. true positives) while triangular leaves represent sequences that do not bind PTx (false positives). Dendrogram A shows the probe which had the most “clonotype-only” predictions, of which 70% are true positives; dendrogram B shows the probe with the most “paratype-only” predictions, of which 75% are true positives; dendrogram C is the probe antibody which is associated with the most false positives (33% true positives) while dendrogram D is the probe antibody associated with the most true positives (20 true positives). Performance is heterogeneous across probes ranging between 33% and 100% precision; precision and recall values reported else-where in this manuscript consider the performance in aggregate of all 364 probes. V and J genes are annotated where this changes within a dendrogram. Dendrograms are constructed using the full VH sequence with the neighbour-joining algorithm of the R package ape [33], plotted using the R package ggtree [34]. The units of the scale bar are amino acid substitions per residue.

Paratyping-only predictions are PTx-binding heavy chain sequences from different clonotypes to the probe antibody. There are 89 predictions which can be split into three groups: those with different V genes (4), those with different J genes (38) and those with CDRH3 identity below 72.0% (47). For example, a pair of binders with just 40% H3 sequence identity but 80% predicted paratope identity were clustered together. Paratype predictions can potentially be further validated by building homology models of the sequences [29]. In this case, modelling of the sequences suggests that the pair of sequences would also be predicted to be highly structurally similar (see Supplementary Table 1 for model information) [35].

### 2.3 Paratyping enables the identification of novel anti-PTx antibodies from different clonotypes to known binders in prospective experiment

#### 2.3.1 Paratyping and clonotyping identify different binders in a bulk repertoire experiment

In order to test how the methods would scale in a larger and non-enriched data set, we performed a prospective repertoire mining experiment in which the heavy chain sequences from PTx-binding antibodies were used to identify novel PTx-binding heavy chains from a set of bulk heavy chain repertoires. We first looked at how the methods scale in terms of number of predictions, and then experimentally validated a number of the predictions of both clonotyping and paratyping.

Using the 364 known PTx binders as probes, 59,107 heavy chain sequences from bulk sequencing repertoires were searched for paratype- or clonotype-related sequences. 4269 sequences were identified by both clonotyping and paratyping, 1113 by paratype-only and 1077 by clonotype-only.

The 1113 paratype-only predictions can be categorised as those with a different V gene (179), those with a different J gene (396) and those with sub-72% CDRH3 identity (538).

#### 2.3.2 Prospective experimental validation of novel PTx-binding sequences

For the experimental validation of predicted PTx binders and non-binders, we created a category of prediction more stringent than the “paratype-only” or “clonotype-only” categories to show the utility of paratyping in a real antibody discovery experiment context where multiple probe sequences are available. As shown in Figure 3, a particular probe antibody may make predictions via clonotype or paratype alone. As reflected in the similarity in the precision-recall values calculated over the aggregate of probe antibodies in the single-cell one-versus-all cross-validation, these “method-unique” predictions become rarer when considering a larger number of probes - such predictions, which could not be found by another method even when using the full complement of known binders, are referred to as “paratype-unique” or “clonotype-unique” predictions.

Of the potential 2193 predicted PTx binders, 139 were selected for expression (see Methods) and PTx-binding assay. An additional 48 antibodies predicted to not bind PTx (due to paratope identity or shared lineage with confirmed non-binders) were also tested. The predictions were split into thirds according to whether they were predicted by both methods (labelled “both”), or were unique across all binders for paratyping (“paratype-unique”) or clonotyping (“clonotype-unique”).

39 of the 43 (90%) novel heavy chains predicted to bind PTx by both paratyping and clonotyping were true PTx binders. Paratope identity between known and predicted binders ranged between 83% and 100%, with CDRH3 identity ranging between 77% and 100%. 31 of the 48 (65%) of the clonotype-unique predictions were true PTx binders, with minimally 57% predicted paratope identity to a known binder, 72% CDRH3 identity and 74% CDRH identity (amino acid identity calculated over the heavy chain CDRs). 14 of the 48 (30%) of the paratype-unique PTx binders bound PTx. The minimal CDRH3 identity of a PTx binder to any known binder was 56% with 76% paratope identity. The distribution of CDRH3, total CDR and total VH amino acid identity of novel PTx-binding heavy chains to known PTx-binding antibodies is shown in figure 5. None of the 48 predicted PTx non-binders bound PTx.

**Figure 4:**
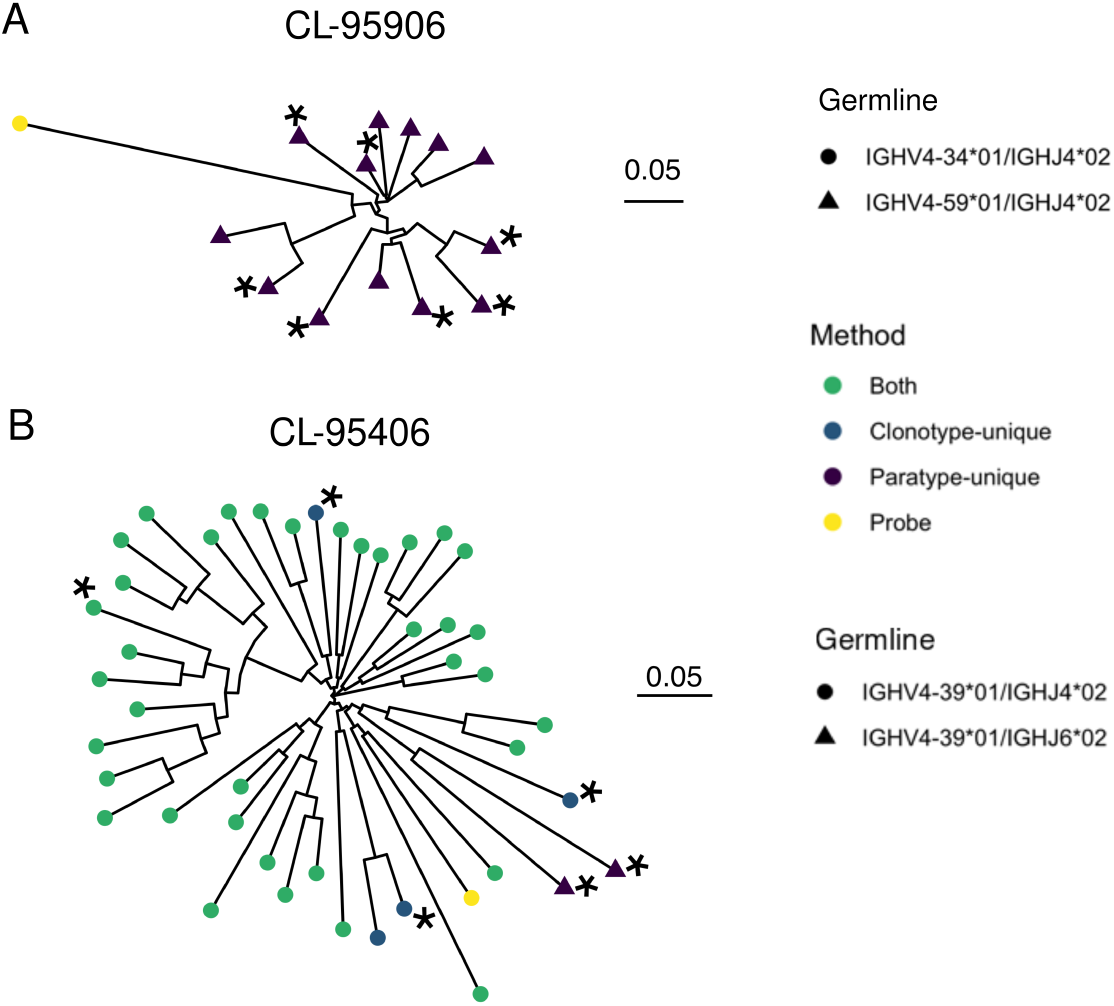
Dendrograms showing two examples of immune repertoire mining, using known PTx binders CL-95906 and CL-95940 (leaves coloured yellow). Heavy chains predicted to bind PTx are coloured as green, blue or purple depending on whether they are predicted to bind via both methods or were clonotype- or paratype-unique. Asterisks indicate heavy chains selected for testing, all of which were validated as PTx binding. A shows sequences predicted to bind PTx that use a different V gene (V4-59) than the known PTx binder used for prediction (V4-34). Sequences using a different J gene to the known PTx binder are shown in B.

**Figure 5:**
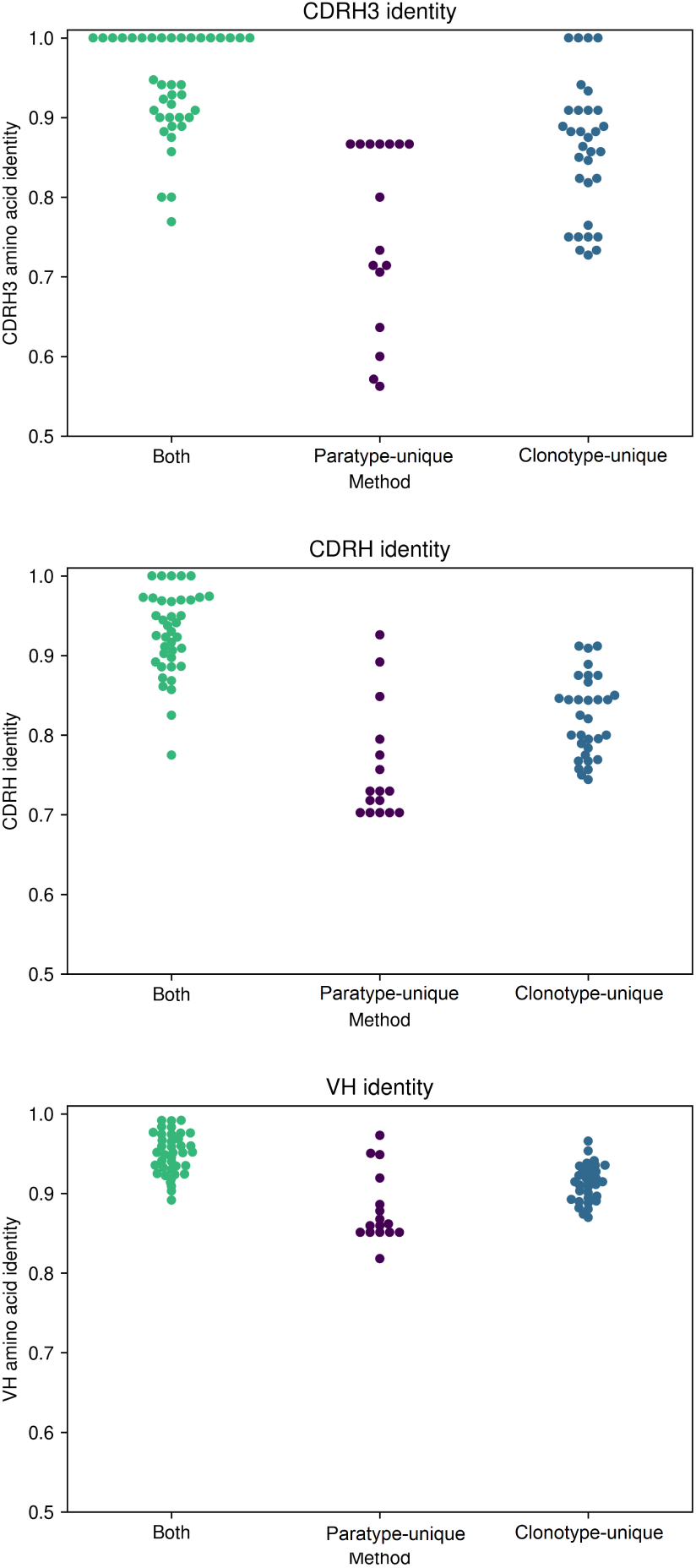
CDRH3, total CDR and total VH amino acid identity of novel PTx-binding antibodies to the known PTx-binding antibody by which they were identified, according to method by which they were identified. Paratyping enables the discovery of PTx-binding sequences with lower sequence identity across each of these regions with minimally 56% CDRH3 identity, 70% CDRH identity and 80% total VH identity.

As in the single-cell data set, the success rate in the predictions which were made by both clonotyping and paratyping is higher than either method alone. The success rate of paratype-unique predictions is significantly lower than that of clonotype-unique predictions. However, it may not be appropriate to compare performance across different probe antibodies, some of which may be liable to activity cliffs (a concept from small molecule chemistry where a compound exhibits a large change in activity given only a small change in structure [36]). A direct comparison can be made where both clonotype-unique and paratype-unique predictions were made using the same known PTx binder. This occurred for 6 PTx probes, and across these an average precision of 75% for paratype-unique and 92% for clonotype-unique was observed.

#### 2.3.3 Discovery of novel anti-PTx antibodies from different clonotypes

Paratyping identified PTx-binding antibodies that could not be found using clonotyping (“paratype-unique”), for example those using a different V gene to any of the known PTx binders. An example is shown in Figure 4), where the original antibody used the inherently autoreactive V gene V4-34 [37], which may be problematic in development. However, paratyping recovers seven PTx-reactive anti-bodies which use the V4-59 gene segment instead.

Paratyping also recovered novel PTx-binding heavy chains which derive from different J genes and examples with CDRH3 identities well below most clonotyping thresholds (commonly 80% - 100%). The minimal CDRH3 identity of a validated PTx-binding antibody to any known binder was 56%, suggesting that paratyping can identify antibodies that bind to the same epitope that could not be found by any clonotyping method.

### 2.4 Repertoire mining can improve in-silico developability metrics

One of the limitations of clonotyping as a method for immune repertoire mining is the relatively narrow sequence space it is capable of making predictions within, meaning that the discovered antibodies may have conserved developability problems. We have already seen one example where paratyping’s ability to jump between germlines allows us to avoid an autoreactive V gene-derived antibody; other developability problems such as aggregation propensity may also be improved by using paratyping.

Of the original 97 antibodies used for repertoire mining, 38 antibodies were flagged by the Therapeutic Antibody Profiler (TAP) tool as having possible developability issues due to CDR length, high density of charge or hydrophobicity, or charge asymmetry between the heavy and light chains. As paratyping only groups antibodies with the same length CDRs, we consider only the latter four developability metrics. Twenty-six of the original probe antibodies were flagged with extreme (2) or unobserved (24) values. Values are considered extreme (flagged as amber by TAP) if they fall within either the top or bottom 5% of the distribution of values observed in a set of 377 clinical-stage therapeutics (CSTs) [38]. Values are unobserved (flagged as red by TAP) if they are outside of the range of values observed in existing CSTs. Seventeen of these probes were used in the identification of novel antibodies assayed for PTx binding, of which thirteen successfully identified new PTx binders. For four of these probes, one of the new PTx-binding antibodies identified showed a sufficiently large change in the flagged developability metric that the flag was removed. There were a further five new PTx-binding antibodies showing a shift in the flagged developability metric towards the mean value among clinical-stage therapeutics (CSTs).

Figure 6 shows the improvement in patch surface hydrophobicity achieved by immune repertoire mining using CL-95375 as a probe antibody. CL-95375 had an amber flag for this metric. Repertoire mining was used to identify a number of predicted PTx-binding antibodies. It can be seen that the more sequence-distinct paratype-only predictions are able to achieve greater changes in PSH. These predictions were not assayed as they were within the clonotype of another known binder and therefore not “paratype-unique".

**Figure 6:**
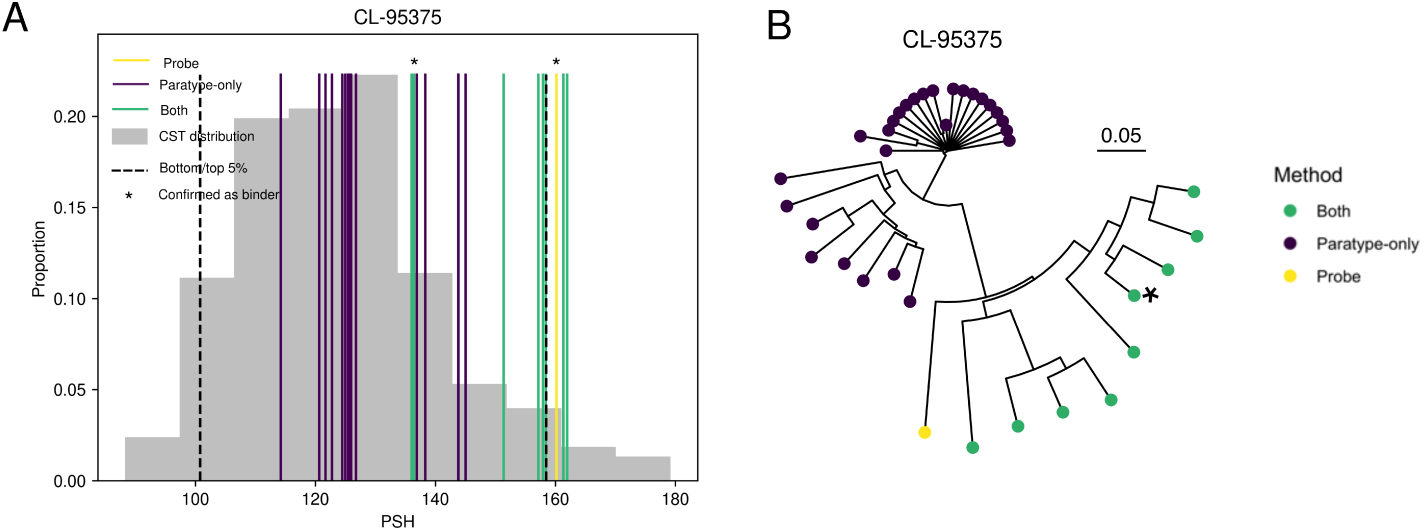
An example of a probe antibody with a flagged developability issue (in this instance, patch surface hydrophobicity (PSH)) which could be significantly improved by immune repertoire mining. Clonotyping and paratyping were used to discover a novel antibody targeting the same epitope but without a developability flag. This shift is shown in A, where asterisks indicate the probe antibody, CL-95375 (yellow), and the validated novel PTx-binding antibody which has a shift in PSH towards the mean value among 277 clinical-stage therapeutics (CSTs) [38]. The extremes of the distribution (upper and lower 5% for PSH) are shown as the black dashed line. The distribution of the metric among antibodies within the same paratype and/or clonotype (colored according to the legend) is shown; the only antibody assayed, which was validated as PTx-binding, is marked with an asterisk. Other examples can be seen in Supplementary Figure 2. The predicted PTx binders in A are shown in the dendrogram in B, with the confirmed novel PTx binder marked by an asterisk.

## 3 Discussion

Characterising the functional relationship between sequence-distinct antibodies is an important step in our understanding of the adaptive immune landscape. Mapping antigen preference to antibody repertoires will allow us to identify epitope convergence of antibodies at a large scale. In a test system of transgenic mice immunised with Pertussis toxoid (PTx), we show for the first time that prediction and comparison of paratopes can be used to group antigen-specific antibodies in both an enriched, single-cell data set and non-enriched bulk heavy chain repertoires. We demonstrate the utility of the method in the context of an antibody discovery experiment alongside the conventional approach of clonotyping and discover new anti-PTx antibodies from different clonotypes to any known binders.

We first developed the method in a single-cell data set, where paratyping and clonotyping were able to group PTx-specific antibody heavy chains with high precision (84% and 82% respectively). These results may not map to the bulk sequencing data set given that the sequences derived from both plasma-, memory- and antigen-sorted cells (leading to ca. 30% of sequences being PTx-reactive). To validate the method in a less enriched data set, we performed a prospective experimental test of paratyping in a non-antigen sorted set of bulk heavy chain sequencing repertoires.

In the prospective experimental test, paratyping allowed us to discover new PTx-binding antibodies that we could not have found using clonotyping. These include antibodies that use different germline genes as well as antibodies with lower CDRH3 identity than in common definitions (80-100%). In terms of antibody discovery, paratyping allows us to identify sequence-distinct antibodies that bind to the same epitope and which can differ significantly in developability or affinity. In the terms of repertoire analysis, paratyping expands our ability to functionally group antibodies beyond clonotypes, and therefore allows us to detect specific cases of epitope convergence between lineages - we found epitope convergence between V4-34/J6 and V4-59/J6 lineages, V4-39/J6 and V4-39/J4 lineages, V4-34/J5 and V4-34/J4 lineages and V5-51/J4 and V5-51/J6 lineages in Pertussis toxoid binders. We did not observe pairs of antibodies in the same paratype using different V gene families. However, paratope identity across the germline-encoded CDRHs 1 and 2 is equal to or in excess of 75% across members of V1 and V7, and V3 and V4 (see Supplementary Figure 3), so we predict that it should be possible to use paratyping to find binders from different V gene families, should a large enough sequencing data set be mined. The implications of this for immune repertoire clustering are as yet unexplored. If such convergence is widespread, it is possible that clonotyping overestimates functional diversity and this may account for low proportions of clonotypes shared across individuals [3].

Success rate in the prospective experimental validation was variable across the categories of prediction. We found that 90% of sequences predicted by both clonotyping and paratyping to be binders bound PTx. The success rate was considerably lower in clonotype-unique predictions (65%) and even lower in paratype-unique predictions (30%). It should be noted that these method-unique predictions form the minority of predictions for either method (15-22% of predictions). The lower success rate of paratyping versus clonotyping may be attributable to the particularly low CDRH3 identity of these predictions to known binders - paratyping does not give special weight to the CDRH3 in the paratope identity calculation despite the particular role it plays in antigen complimentarity. Predictions with as little as 33% CDRH3 identity to a known binder were assayed but no antibody with an H3 amino acid identity below 56% bound to Pertussis toxoid, suggesting that paratyping could be further improved by the use of CDRH3 weighting.

Paratope prediction only takes around 0.1 seconds per sequence (of which 0.02 seconds corresponds to CDR extraction) as opposed to 0.05 seconds per sequence for VDJ annotation using IgBlast [39]) which means it is considerably more tractable for large datasets than homology modelling (on average 30 seconds per sequence, 600 times slower than germline gene annotation [29]). It has the advantage that it does not rely on the upkeep of consistent and complete germline databases (a leading cause of disagreement between germline annotation tools [40]), but rather on the distribution of a pretrained paratope prediction model [41]. The paratope prediction step is purely sequence-based and structural modelling is not required, meaning that immune repertoires can be annotated without accurate germline alignment and without access to large compute power.

Paratyping was shown to identify functional relationships between PTx-binding antibodies that are not related by clonotype; we would expect this to generalise across protein antigens. As an example, we looked at Cov-AbDab [42], a database of antibodies and nanobodies known to bind to betacoronavirus proteins. Paratyping identified a number of pairs of antibodies from different clonotypes binding to the same epitope, for example MERS-4 and MERS-4-V2, D12 and MERS-27, and 27D and 6A. D12 and 4C2, shown in Figure 1, are another example.

Deciphering the functional landscape of immune repertoires will greatly improve our understanding of the adaptive immune system. Improving our ability to group antibodies binding to the same epitope is a step towards this. Our results show here that the simple and computationally rapid abstraction of the antibody binding site used by paratyping is sufficient to group antigen-specific antibodies in a way which provides us with additional information to clonotyping. This additional information is particularly significant in the context of antibody discovery, where it allows us to recover different novel antigen-specific antibodies from immune repertoires.

## 4 Methods and Materials

### 4.1 Data sets

#### 4.1.1 Single-cell data set

Five genetically engineered mice that have a full set of human immunoglobulin variable region genes

[31] were immunised with Pertussis toxoid (PTx). 1290 paired (*V*_*H*_ /*V*_*L*_) sequences were recovered from antigen-sorted, plasma and memory cells via a previously published method [43].

These 1290 sequences were expressed in HEK293 cells. Antibody supernatant was collected on day eight after transfection and screened for binding to wildtype PTx by homogeneous time-resolved fluorescence (HTRF). Positive control anti-PTx antibody (ab37574, abcam) was diluted in Expi293™ Expression Medium (Gibco) over an 11-point titration using 1 in 3 dilutions to generate a standard curve. Titrations of 5 l of antibody (ab37574) were added to a 384-well white walled assay plate (Greiner Bio-One). Negative control wells received 5 l of Expi293™ Expression Medium only. 5 l of HEK293 antibody supernatants (undetermined concentration) were added to one well of the 384-well plate. 5 l of PT conjugated to Alexa 647 (Lightning-Link, Innova Bioscience) (3.75 nM final concentration) was added to all wells of the assay plate except negative control wells, which instead received 5 l Expi293™ Expression Medium. Finally, 10 l of anti-mouse IgG donor antibody (Southern Biotech; to bind to murine constant region in control and chimeric Kymouse antibodies) labelled with europium cryptate (Cis Bio), (1:4000 final concentration) was added to each well and the assay was left in the dark at RT to incubate for 2 h. After incubation, the assay was read on an Envision plate reader (Perkin Elmer) using a standard HTRF protocol. 620 nm and 665 nm channel values were exported to Microsoft Excel (Microsoft) and F calculations performed ((665/620nm ratio – signal negative control)/signal negative control) * 100). Percent effect values were calculated for each antibody by comparing its F value against a positive control antibody (ab37574) at 6.66 nM.

Antibody sequences with greater than 10% effect value relative to the positive control were labelled as binders. This resulted in 364 PTx-binders and 926 non-binders.

#### 4.1.2 Bulk data set

Heavy chains from sorted splenic B-cells from the same individual five mice as the single-cell data set were sequenced using standard protocols [20] and processed using the pRESTO/Change-O pipeline [44, 45]. This resulted in 259,151 heavy chain sequences. For quality control, only sequences with read count (reads with a particular unique molecular identifier (UMI)) above or equal to two or a consensus count (reads with different UMIs but the same nucleotide sequence) above or equal to ten were considered, reducing the size of the data set to 59,107 sequences.

### 4.2 Clonotyping

Clonotypes were defined as groups of heavy chain sequences sharing the same V and J genes, with identical CDRH3 lengths and a number of amino acid mismatches equal to or below a threshold sequence identity. VJ annotation was performed with IgBLAST [39] within Change-O [45]. CDRH3s were extracted according to the North definition [46] with IMGT numbering [47] performed with ANARCI [12].

### 4.3 Paratyping

Paratypes were defined as heavy chain sequences sharing the same CDR lengths and greater than a threshold sequence identity across the predicted paratope regions. CDRs were extracted according to North definitions [46] with IMGT numbering [47] performed via ANARCI [12].

Parapred [41] was used for paratope prediction using the model as distributed by E. Liberis at https://github.com/eliberis/parapred. To convert the output of Parapred, binding probabilities, into a binary label, we selected a threshold of 0.67 as deemed optimal by the authors of the original paper [41], i.e. residues with a predicted probability of being in the paratope of above 0.67 were annotated as paratope residues. Paratope identity was defined as the number of identical paratope residues (residues which are predicted to be in the paratope in both cases) divided by the smallest number of paratope residues of either sequence being compared (figure 7).

**Figure 7:**
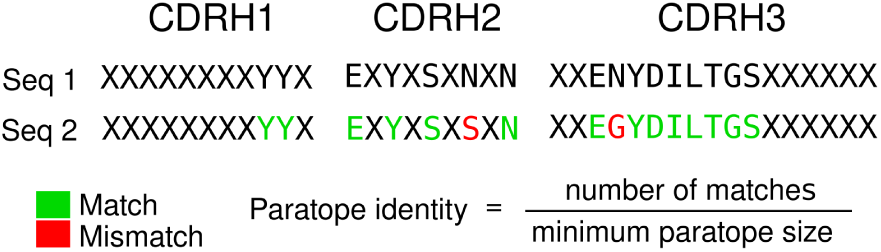
Method of calculating predicted paratope identity. X indicates CDR residues not predicted to constitute the paratope. In the example shown, the paratope identity is 87.5%.

### 4.4 Repertoire mining experiment

In the repertoire mining experiment, we selected a number of hit antibodies from the single-cell data set to use as probes. SPR was carried out on the HTRF-positive antibodies and the highest affinity representatives were selected from each clonotype containing only sequences labelled as binders equating to 97 antibodies, in order to align with previous repertoire mining experiments [9]. We also selected a number of non-PTx binding antibodies to be used as a negative control. The non-PTx binding antibodies were selected as representatives of clonotypes containing only non-PTx binding sequences (551 antibodies).

The heavy chains from these probe antibodies were used as probes to mine the bulk repertoires via both paratyping and clonotyping, using the optimal sequence identity thresholds from the single-cell data set (75% predicted paratope identity for paratyping and 72% CDRH3 sequence identity for clonotyping). The paratyping process is illustrated in Figure 8.

**Figure 8:**
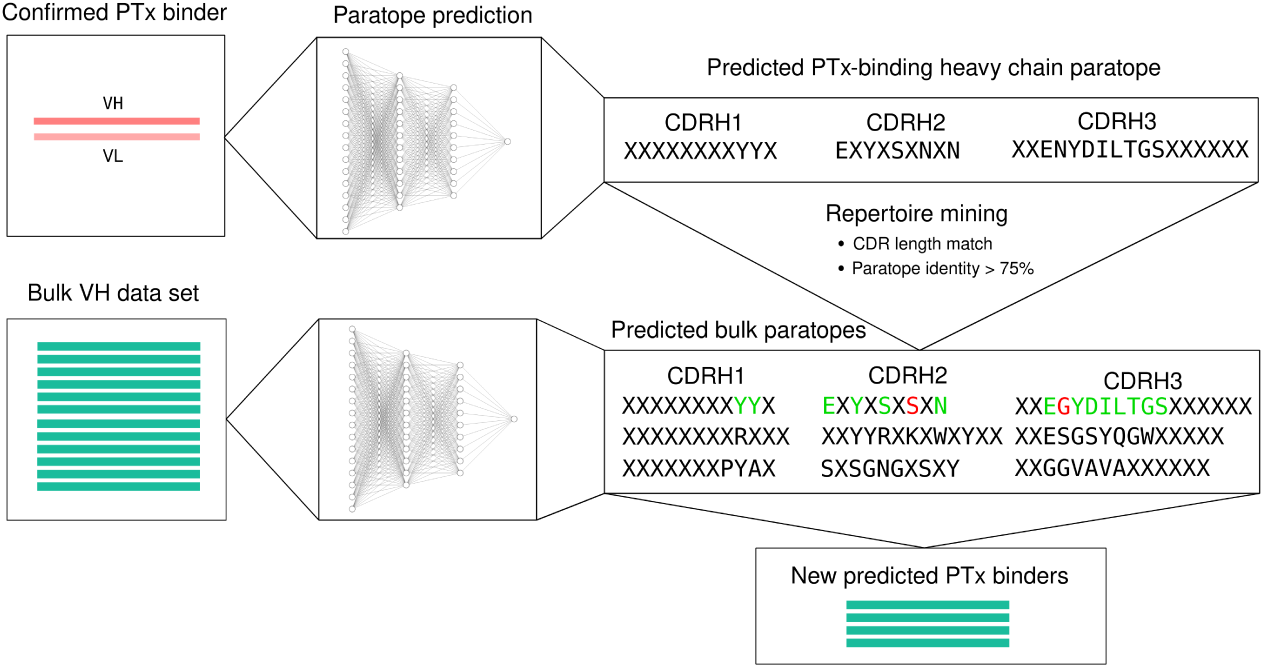
Graphical illustration of the process of repertoire mining via paratyping. A probe antibody is selected. Paratope prediction is performed both on the probe antibody and the bulk data set. Heavy chain predicted paratope identity is used to mine the bulk repertoire for new predicted binders.

Predictions were labelled as “paratype-only", “clonotype-only” or “both” as detailed in the Results section. If an antibody is within the same paratype as a particular probe but not within its clonotype, it is a “paratype-only” prediction and vice-versa. A prediction is labelled as “both” if it is within both the paratype and clonotype of the same probe. In the “Experimental validation of predicted binders” section of Methods, the more stringent “paratype-unique” and “clonotype-unique” definitions were used.

#### 4.4.1 Experimental validation of predicted binders

We selected 192 heavy chain sequences from the bulk data set for expression. 144 of the sequences were predicted to be PTx binders using the 97 PTx-binding probe sequences. We created a new way of categorising predictions in order to fully test the method in the context of an antibody discovery experiment. We define “paratype-unique” predictions as predictions which are not within the paratype of not only the probe in question (paratype-only) but of any of the full complement of 97 probes, and similarly for “clonotype-unique” predictions. Paratype-unique predictions are a more stringent subset of paratype-only predictions and present a greater challenge of the method (as such predictions tend to be corroborated by fewer probes). 48 predicted PTx-binding heavy chain sequences were selected from each of the three categories of prediction (“paratype-unique", “clonotype-unique” and “both”). Five of the “both” predictions were identical to their probes. The remaining 48 sequences assayed were predicted non-PTx binding heavy chain sequences with 16 sequences selected from “clonotype-unique", “paratype-unique” and “both” categories of prediction.

The heavy chain sequences selected for expression from the bulk repertoire were paired with the cognate light chain of the probe sequence by which they were identified. Within-clonotype VH/VL pair-ing has been validated [9, 48] as a method of reconstituting binding where the cognate light chain is not sequenced. Pairings were only made where there was greater than 82% sequence identity across the residues considered to constitute the *V*_*H*_ /*V*_*L*_ interface, based on solvent accessibility [38], in order to maximise the probability of expression. These positions lie outside of the CDRs and therefore do not enforce any constraint on binding site sequence identity.

The predicted binding and non-binding antibodies were expressed in HEK293 cells. The assay is as described in the single-cell data set section of Methods with the exception that the anti-PTx antibody 1B7 was used as positive control. Antibody sequences with an F value exceeding 100 were labelled as PTx-reactive antibodies.

### 4.5 Performance evaluation

To evaluate the performance of either method in grouping PTx-binding sequences, we calculated precision and recall of the method in the single-cell data set according to the following definitions of true positives, false positives and false negatives.

The task is to group antibodies that bind Pertussis toxoid. However, the method is hypothesised to work by grouping by epitope. There will be multiple epitopes on Pertussis toxoid so not all PTx-binding antibodies will be grouped into a single paratype or clonotype. As a result, we do not expect perfect recall. Further, we do not classify antibodies as non-binders if they do not group with a particular binder-we only predict that they do not bind at the same epitope as the binder in question. As a result, we do not calculate a “true negative” rate.

- True positive (TP): A PTx-binding sequence that was identified by another PTx-binding sequence.
- False positive (FP): A non-PTx-binding sequence that was identified by a PTx-binding sequence.
- False negative (FN): A PTx-binding sequence that was not identified by any PTx-binding sequence.

### 4.6 In-silico developability assessment

We calculated the in-silico developability metrics of total CDR length, patches of surface hydrophobicity (PSH), patches of positive charge (PPC), patches of negative charge (PNC) and structural Fv charge symmetry parameter (SFvCSP) using the Therapeutic Antibody Profiler tool (TAP) [38]. The tool calculates these metrics from homology models built using ABodyBuilder [35] and compares them to a database of 277 clinical-stage therapeutics (CSTs) [49]. According to the metric in question, values are flagged as “amber” (extreme) if they lie outside the upper or lower, or upper and lower, 5% of observed values among CSTs. An antibody is flagged as “red” if the value of a particular metric is outside of the observed range in CSTs. [38]

## 6 Data and code availability

The code and data associated with this study are available at http://opig.stats.ox.ac.uk/resources.

