## Supplementary Information for "A computational method for immune repertoire mining that identifies novel binders from different clonotypes, demonstrated by identifying anti-Pertussis toxoid antibodies"

Eve Richardson[1], Jacob D. Galson[2,3], Paul Kellam[4,5], Dominic F. Kelly[6,7], Sarah E. Smith[4], Anne Palser[4], Simon Watson[4], and Charlotte M. Deane*[1]

[1]Department of Statistics, University of Oxford, UK
[2]Alchemab Therapeutics Ltd, London, UK
[3]University Children's Hospital, University of Zurich, Switzerland
[4]Kymab Ltd, Cambridge, UK
[5]Department of Infectious Disease, University College London, UK
[6]Department of Paediatrics, University of Oxford, UK
[7]Oxford University Hospitals NHS Foundation Trust, Oxford, UK

| ID | Framework | CDRH1 | CDRH2 | CDRH3 |
|---|---|---|---|---|
| **CL-97155** | 3k2u | 5uea | 6elu | 1iqd |
| **CL-97116** | 3k2u | 5uea | 6elu | 1iqd |

Table 1: PDB IDs of templates used in homology modelling of a pair of antibodies with just 40% CDRH3 amino acid identity but 80% predicted paratope identity, as referred to in the 'Paratyping and clonotyping successfully cluster PTx binders in a single-cell dataset' section of Results. Sequences were homology modelled using ABodyBuilder [2] which assigns structural templates to each of the CDRs and the framework region. Common template usage across all of the CDRs and framework region indicates structural similarity (structural similarity despite low sequence identity is noted elsewhere [3]). The relationship between these two Pertussis-toxoid binding antibodies was recovered through paratyping, and demonstrates that low sequence identity hits can have both chemically and structurally similar binding sites.

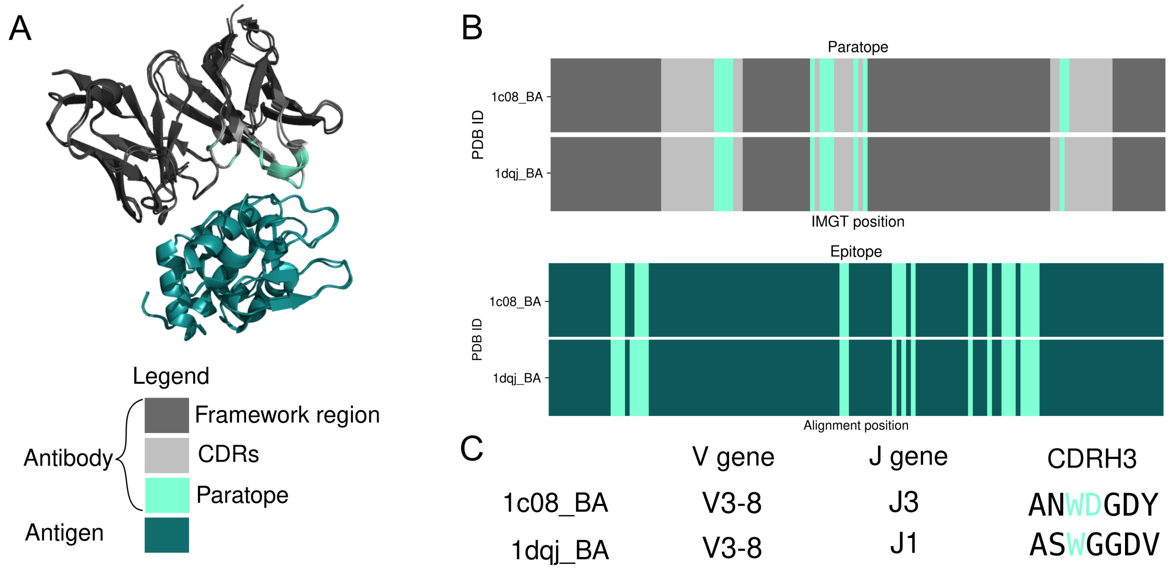


Figure 1: An example of two antibodies that bind to the same epitope but derive from different clonal lineages. HyHel-10 (PDB ID: 1c08) and HyHel-63 (PDB ID: 1dqj) anti-hen egg white lysozyme antibodies target the same residues, despite being derived from different J genes and displaying CDRH3 amino acid identity (57.1%) below the standard clonotyping definition (80% - 100%) (C). The antibodies use the same paratope residues (100% paratope identity) (B) to achieve this functional convergence (95.7% epitope identity). The paratope and epitope are defined as those residues with any atom within 4.5 Å of any residue in the antigen or antibody respectively. Epitope and paratope identity correspond to sequence identity at equivalent epitope and paratope residues in the antigen and antibody alignments respectively, where the denominator is the minimum number of epitope or paratope residues in the pair of structures being compared.

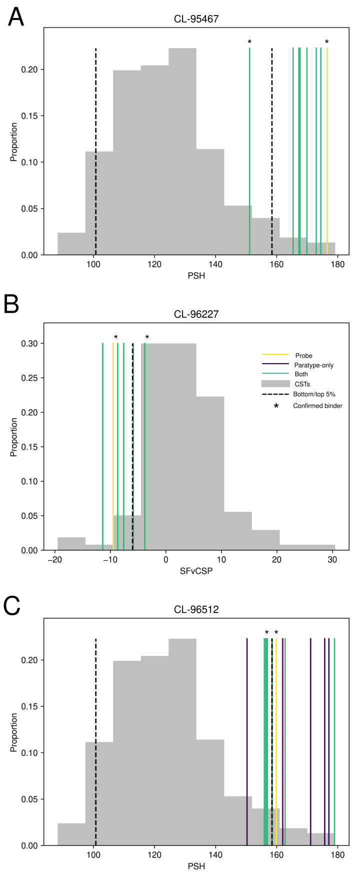


Figure 2: As referred to in the 'Repertoire mining can improve in-silico developability metrics' section of Results, three further examples of probe antibodies with flagged developability issues (in figures A and C, patch surface hydrophobicity (PSH); in figure B, charge asymmetry between the heavy and light chains (SFvCSP)) where the developability flag was removed in putative PTx binders identified by immune repertoire mining. Asterisks indicate confirmed PTx-binding antibodies, the probe antibody (yellow) and binders identified through prospective experimentation. The putative PTx-binding antibodies may identified by both clonotyping and paratyping (green), paratyping only (purple) or clonotyping only (blue). In these three instances, the change in the property is significant enough that the identified PTx-binding antibody discovered through repertoire mining is no longer flagged. Developability flags are assigned using the Therapeutic Antibody Profiler, TAP [1], which calculates five metrics associated with developability from homology models and compares these values with the distribution observed among clinical-stage therapeutics (CSTs); antibodies that lie in the extremes of a particular metric are "flagged" with respect to that metric.

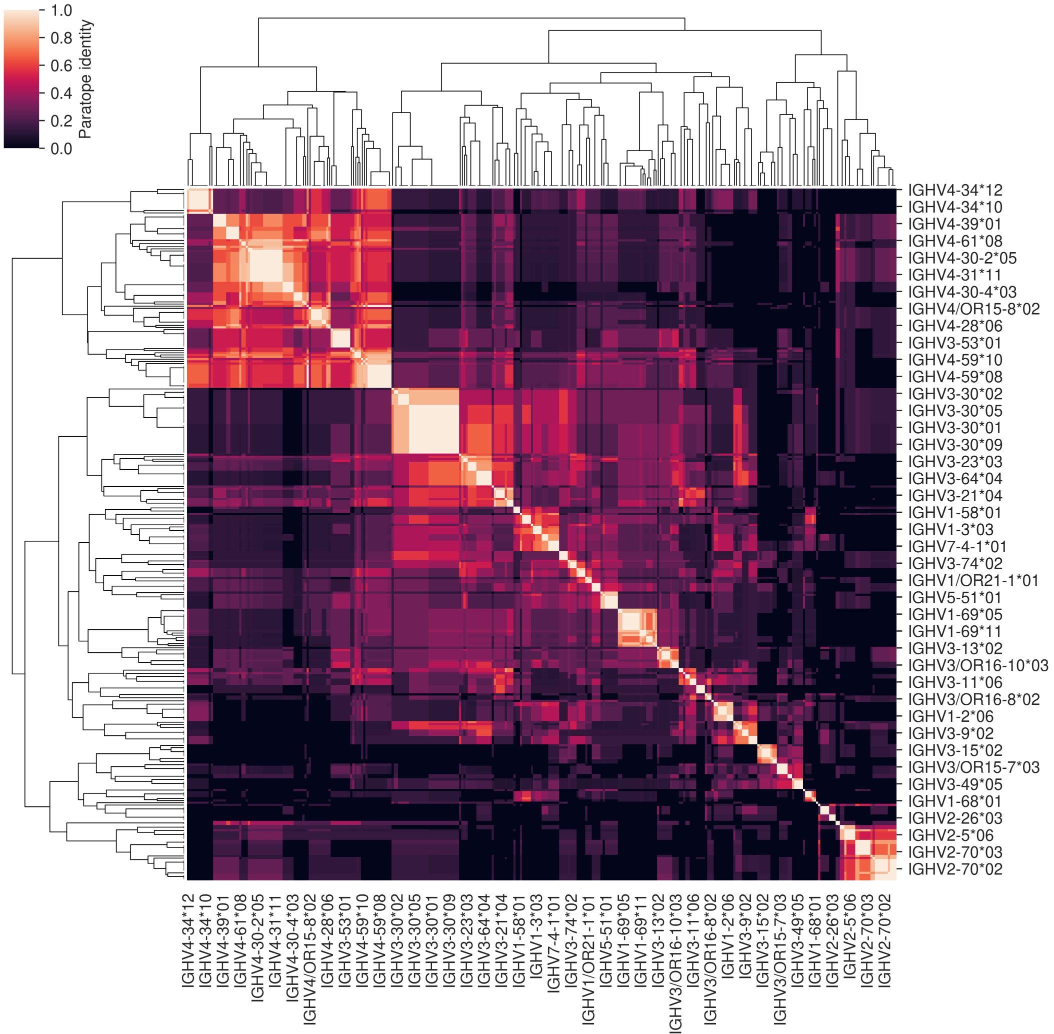


Figure 3: Predicted paratope identity across CDRH1 and CDRH2 of germline alleles (IMGT, as of February 14th 2020). Paratope identity may be equal to or in excess of 75% across V genes within the same and different families - this was detected in alleles from genes IGHV3-53 and IGHV4-59, IGHV3-66 and IGHV4-59 and IGHV7-4-1 and IGHV1-8, suggesting that in large enough sequencing data sets paratyping could cluster antibodies from different V gene families.
